## Supplementary figures for "Charge and hydrophobicity are key features in sequence-trained machine learning models for predicting the biophysical properties of clinical-stage antibodies"

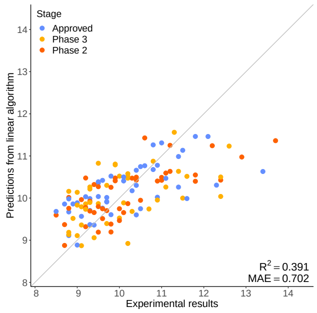


(a) HIC

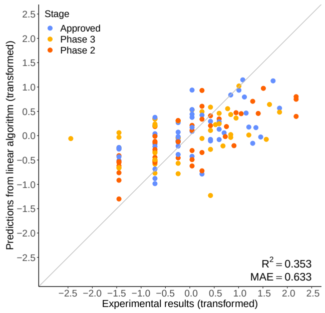


(b) SMAC

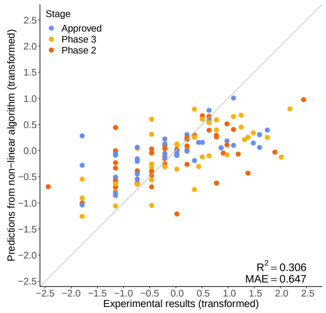


(c) CIC

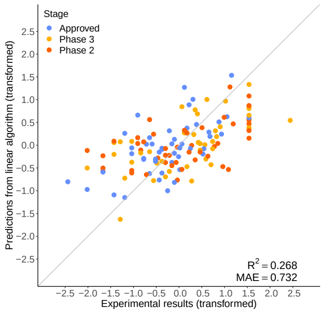


(d) ACSINS

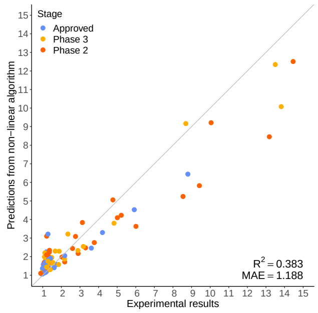


(e) ELISA

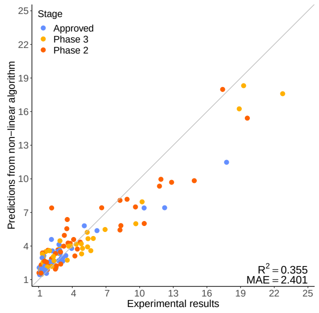


(f) BVP

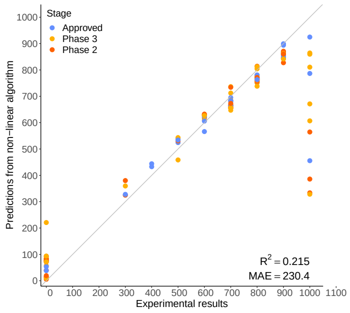


(g) SGAC-SINS

Figure S1: Better performing models. The scatter graphs demonstrate the predictive power of each model, where the original experimental value is on the $x$ axis, and the prediction is on the $y$ axis. The closer each data point is to $y = x$ the better the prediction. For example, in Figure A, the predictions for HIC are generally close to the $y = x$ line until $x$=11 minutes. Above this point the model begins to lose predictive power, and the values begin to fall away from the diagonal. Where applicable, both the $x$ and $y$ axis are reported in terms of the mathematically transformed values.

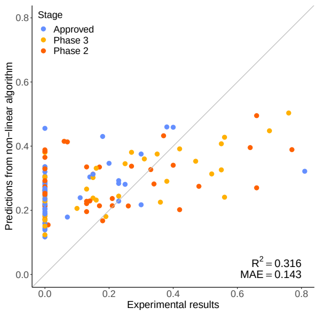


(a) PSR

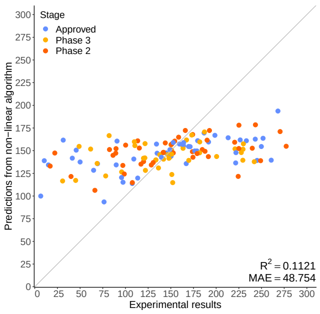


(b) HEK

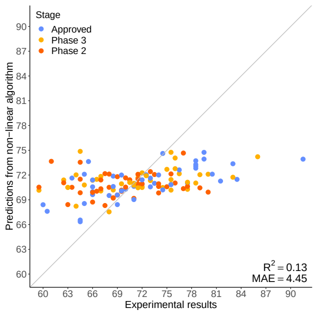


(c) DSF

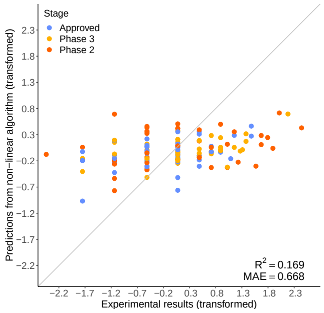


(d) CSI

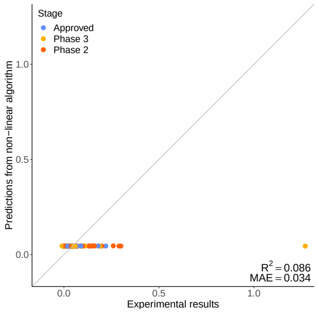


(e) ACC STAB

Figure S2: Less well performing models. As in Figure S1, the scatter graphs demonstrate the predictive power of the models. In comparison, these models do not perform as well and therefore have lower predictive power.

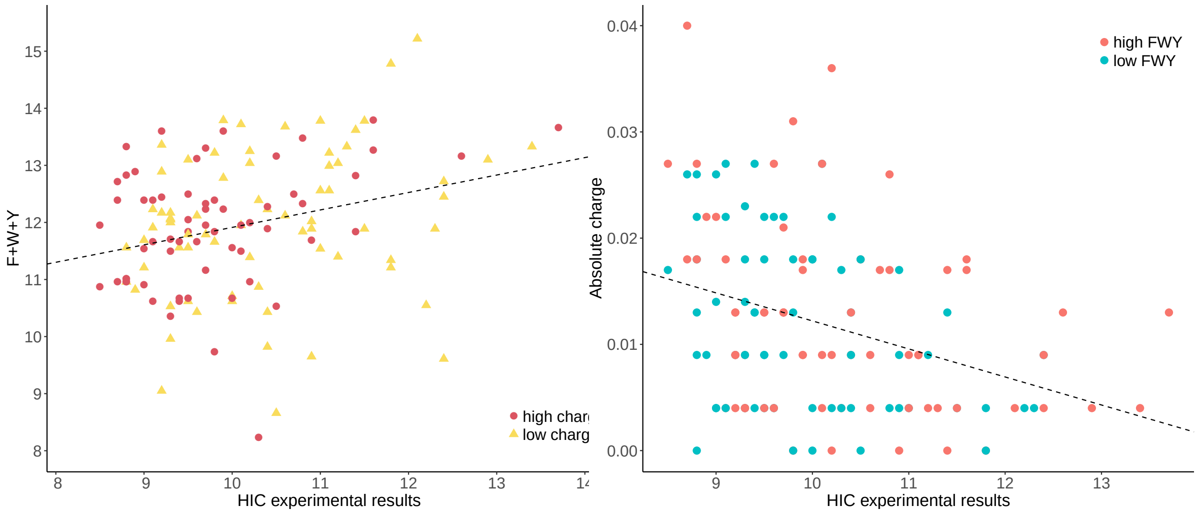


(a) Aromatic content (b) Charge

Figure S3: HIC experimental results for each protein in the mAb137 dataset in context of the aromatic content and absolute charge for each Fv sequence. In A, the aromatic content — here defined as the combined composition value for the amino acids F, W and Y — shows a general positive correlation with the HIC experimental score for that sequence. Sequences with an above average absolute charge value, in the context of the mAb137 dataset, are coloured red, and those with less yellow. In B, there is a general negative correlation between the absolute charge — calculated from the number of charged residues D, E, K and R within the sequence — and HIC experimental score. Sequences with an above average aromatic content are coloured pink, and those with a below average aromatic content are coloured blue.
